# Raw sequence to target gene prediction: An integrated inference pipeline for ChIP-seq and RNA-seq datasets

**DOI:** 10.1101/220152

**Authors:** Nisar Wani, Khalid Raza

## Abstract

Gene expression patterns determine the manner whereby organisms regulate various cellular processes and therefore their organ functions.These patterns do not emerge on their own, but as a result of diverse regulatory factors such as, DNA binding proteins known as transcription factors (TF), chromatin structure and various other environmental factors. TFs play a pivotal role in gene regulation by binding to different locations on the genome and influencing the expression of their target genes. Therefore, predicting target genes and their regulation becomes an important task for understanding mechanisms that control cellular processes governing both healthy and diseased cells.In this paper, we propose an integrated inference pipeline for predicting target genes and their regulatory effects for a specific TF using next-generation data analysis tools.

## 1 Introduction

Omics technologies are key drivers of the data revolution that has taken place in the life sciences domain from last few decades. These technologies enable unbiased investigation of biological systems at genomic scales. Using high throughput Next Generation Sequencing (NGS) methods, genome-wide data is collected from cells, tissues and model organisms (Raza & Ahmad, 2016). These data are key to investigate biological phenomena governing different cellular functions and also help biomedical researchers to better understand the disease etiologies which have not been previously explored. NGS protocols such as, ChIP-seq and RNA-seq are to generate datasets from where we can obtain genome-wide binding map of TFs and epigenetic signatures(Park, 2009; Furey, 2012) and can also measure the gene expression abundance within the cell for the whole genome (Costa, Angelini, De Feis, & Ciccodicola, 2010; Wang, Gerstein, & Snyder, 2009; Ozsolak & Milos, 2011).

Numerous efforts have been put forth to uncover the interplay between genomic datasets obtained from ChIP-seq and RNA-seq for gene regulation studies of individual TFs (Wang. et al., 2013) or mapping Transcription Regulatory Networks as in (Wade, 2015). Revealing such interaction between these data has significant biomedical implications in various pathological states as well as in normal physiological processes (Yue et al., 2014).Therefore, there is a compelling need to integrate these data to predict the pattern of gene expression during cell differentiation (Kadaja et al., 2014) and development (Comes et al., 2013) and to study human diseases such as, cancer as outlined in (Portela & Esteller, 2010).

The aim of this study is to integrate genome-wide protein DNA interaction (ChIP-seq) and transcriptomic data (RNA-seq) using a multi-step bioinformatics pipeline to infer the gene targets of a TF which serve as building blocks of a transcriptional regulatory network.We have developed a Perl script that implements this multistage pipeline by integrating tools in the same order as depicted in Fig. (1). The choice of tools for each stage is a consequence of thorough literature study among the set of tools available in their respective domains.Our implementation is a partially automated system that requires supervision at the time of quality control of raw reads, but progresses smoothly onwards without any manual intervention to integrate the two datasets and generate TF-specific gene targets.

**Fig.1:**
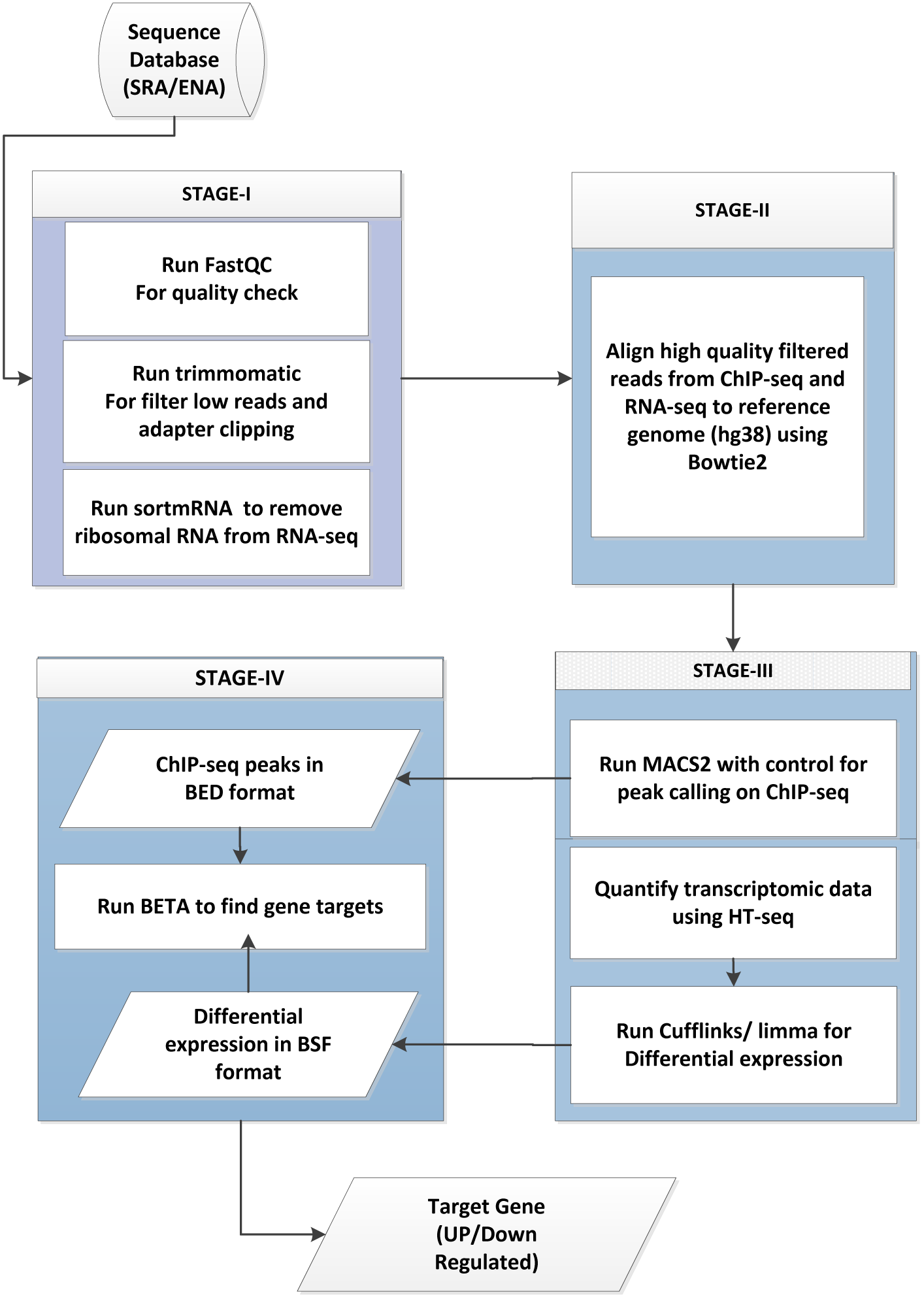
Proposed inference pipeline for target gene prediction

## 2 Related Literature

Software tools and methods exist that predict and analyze gene targets by processing ChIP-seq data. A distinguishable group of peak callers such as, CisGenome (H. Jiang, Wang, Dyer, & Wong, 2010), BayesPeak (Spyrou, Stark, Lynch, & Tavaré, 2009), Model-based Analysis of ChIP-seq (MACS) (Zhang et al., 2008),Peakseq (Rozowsky et al., 2009), SICER (Zang et al., 2009) are some of the widely used tools that identify TF-binding sites. These peak callers identify the target genes either by looking for peaks in promoter region or assign a proximal nearest gene in the vicinity of peaks. However, with most TFs ChIP-seq data having peaks in and around the promoter regions is very less. Also predicting targets using nearest peak is not always reliable.TIP (Cheng, Min, & Gerstein, 2011) is another tool that builds a probabilistic model to predict gene targets, but does not take into account gene expression data. Certain databases such as JASPAR (Sandelin, Alkema, Engström, Wasserman, & Lenhard, 2004), TRED (C. Jiang, Xuan, Zhao, & Zhang, 2007) etc. identify target genes for a selected set of TFs based on the motif analysis of the promoter regions using Position Weight Matrices (PWMs). A recent study (Essebier, Lamprecht, Piper, & Boden, 2017) combines multiple approaches to predict target genes.

On the contrary, some earlier studies used gene expression data for predicting target genes. (Qian et al., 2003) use Support Vector Machines (SVM) to discover relationships between the TFs and their targets; (Honkela et al., 2010) identify targets with time-series expression data by creating a linear activation model based on Gaussian process.

## 3 Materials and Methods

The proposed pipeline operates on raw NGS data, ChIP-seq & RNA-seq. After preprocessing the raw sequences it yields differential gene expression and peak information of genes from these data sets. Both these datasets are integrated to yield target genes for the ChIPred TF.A working description of the proposed pipeline is presented below.

### 3.1 Datasets

NGS data is primarily accessed from Sequence Read Archive (SRA) at NCBI (Leinonen. & Sugawara., 2010) and European Nucleotide Archive (ENA) at EBI (Leinonen et al., 2010). Raw sequences in the form of FASTQ files are freely available for download for a variety of cell types, diseases, treatments and conditions. Besides the public databases, a number of projects and consortia offer public access to their data repositories.For example, ENCODE(Consortium et al., 2004) is a publicly funded project that has generated large sets of data for a variety of cell lines, tissues and organs. Raw as well as pre-processed data can be accessed and freely downloaded from ENCODE data portal. Another publicly funded research project, The Cancer Genome Atlas (TCGA) (Weinstein et al., 2013) also provides datasets for a variety of cancer types. For the current study we have downloaded ChIP-seq data of MCF7 breast cancer cell line from ENCODE experiment *ENCFF580EKN* and RNA-Seq data of transcriptomic study *PRJNA312817* from Eurpean Nucleotide Archive. The RNA-seq experiment contains 30 samples of time course gene expression data from MCF7 cell line subjected to estrogen stimulation.

### 3.2 Pipeline Workflow

Molecular measurements within the NGS data exist in the form of millions of reads and are stored as FASTQ files. Information within the raw files is hardly of any value and needs extensive pre-processing before this data can be analyzed. The pre-processing task is a multi-step process and involves the application of a number of software tools. In this section, we present a detailed NGS pipeline that describes necessary steps from pre-processing of RNA-Seq and ChIP-seq data to target genes regulated by ChIPred TF.The pipeline has been implemented using a Perl script by integrating various NGS data processing & analysis tools.Fig. (1) is a graphical depiction of the proposed inference pipeline.

### Quality Control

Almost all sequencing technologies produce their outputs in FSATQ files. FASTQ has emerged as the de facto file format for data exchange between various bioinformatics tools that handle NGS data. FASTQ (Cock, Fields, Goto, Heuer, & Rice, 2009) format is a simple extension to existing FASTA format; the files are plain ASCII text files with the ability to store both nucleotide sequences along with a corresponding quality score for each nucleotide call.

Base sequence qualities are usually interpreted in terms of Phred quality scores. Phred quality scores *Q* are defined as a property which is logarithmically linked to error probabilities *P* of called bases and can be computed as shown in equation (1).

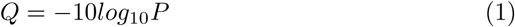

Phred’s error probabilities have been shown to be very accurate (Ewing, Hillier, Wendl, & C., 1998), e.g. if Phred assigns a quality score of 10 means that 1 out of 10 base calls is incorrect, a score of 20 depicts that 1 in 100 bases has been called incorrectly.Usually, a Phred score >= 20 is considered as acceptable read quality, otherwise read quality improvement is required. If read quality is not improved by trimming, filtering, and cropping, there may be some error during library preparation and sequencing. FASTQC (Andrews et al., 2010) is a Java based tool that is used to assess the quality of the reads produced by Next Generation DNA Sequencers. Low quality reads are excised from the FASTQ files to improve the quality of the reads. Various tools are available that can be used to trim bases with poor Phred scores i,e. Phred score less than 20.

Trimmomatic (Bolger, Lohse, & Usadel, 2014) is a Java based open source tool used for trimming illumina FASTQ data and removing adapters. Additionally RNA extracted using NGS does include non-coding RNA molecules besides the coding ones. These non-coding RNAs are usually the ribosomal RNA. For quantifying the gene expression patterns using RNA-seq, it is essential that these non-coding RNAs be filtered from the existing reads. SortMeRNA (Kopylova, Noé, & Touzet, 2012) is a very efficient and accurate tool that is used to filter out the ribosomal RNA from the metatranscriptomic data.A diagrammatic flow of qulaity check and trimming is shown in Fig.(2).

**Fig.2:**
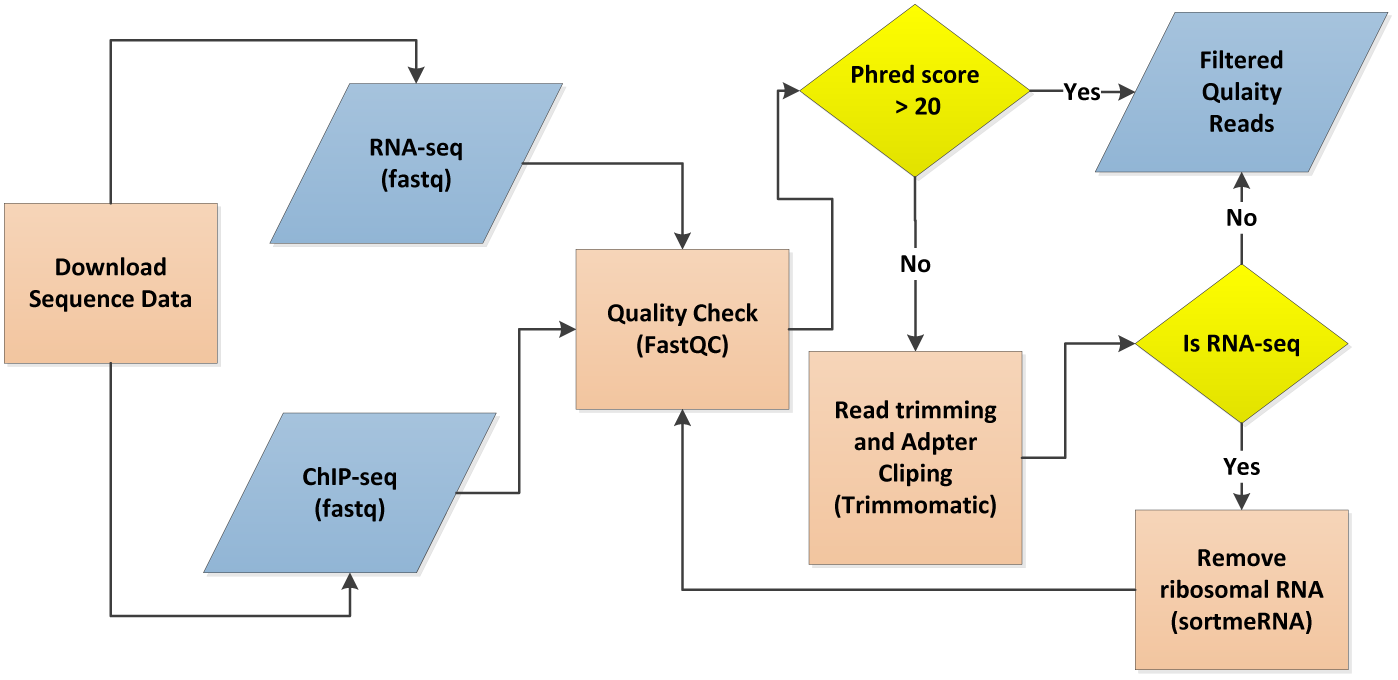
Flowchart showing quality control check, adapter clipping and trimming steps of raw reads

### Mapping sequence reads to a reference genome

With required read quality achieved after trimming and adapter clipping, the next step is to align the short reads to a reference sequence. The reference sequence is our case is human genome assembly hg38/hg19 but it can also be a reference transcriptome, or a de novo assembly incase a reference sequence is not available. There are numerous software tools that have been developed to map reads to a reference sequence. Besides the common goal of mapping these tools vary considerably from each other both in algorithmic implementation and speed. A brief account can be found in (Flicek & Birney, 2009)

In this pipeline we used Bowtie2 genome aligner (Langmead & Salzberg, 2012), because it is a memory-efficient and an ultrafast tool for aligning sequencing reads. Bowtie2 performs optimally with read lengths longer than 50bp or beyond 1000bp (e.g. mammalian) genomes. It builds an FM Index while mapping reads to keep its memory footprint small and for the human genome it is typically around 3.2 GB of RAM.

### Expression quantification of RNA-Seq

Although raw RNA-seq reads do not directly correspond to the gene expression, but we can infer the expression profiles from the sequence coverage or the mapping reads that map to a particular area of the transcriptome. A number of computational tools are available to quantify the gene expression profiles from RNA-seq data. Software tools, such as Cufflinks, HTSeq, IsoEM, and RSEM are freely available. A comparative study of these tools is presented in (Chandramohan, Wu, Phan, & Wang, 2013).

Despite clear advantages over microarrays, there are still certain sources of systematic variations that should be removed from RNA-seq data before performing any downstream analysis. These variations include between sample differences, such as sequencing depth and within sample differences e.g. gene length, GC content etc. In order to circumvent these issues and exploit the advantages offered by RNA-seq technology, the reads/ kilobase of transcript per million mapped reads (RPKM) normalizes a transcript read count by both its length and the total number of reads in the sample (Pepke, Wold, & Mortazavi, 2009). For data that has originated from the paired-end sequencing, a similar normalization metric called FPKM (fragments per kilobase of transcript per million mapped reads) is used. Both RPKM and FPKM use similar operations for normalizing single end and paired end reads (Conesa et al., 2016).

Counts per million (CPM) is another important metric provided by limma package to normalize gene expression data. Once the normalized expression estimates are available, we can obtain a differential expression of gene lists across the samples or conditions using limma voom (Ritchie et al., 2015) provided by R bioconductor.

**Peak Calling** Early pre-processing steps of ChIP-seq data resemble that of RNA-seq. Beginning with the quality check of raw reads, read trimming and adapter elimination, filtered high quality reads are then mapped to reference genome using Bowtie2 as described above.

In order to identify the genomic locations where the protein of Interest (POI) has attached itself to DNA sequences, the aligned reads are subjected to a process known as Peak Calling. Software tools that predict the binding sites where this protein has bound itself by identifying location within the genome with significant number of mapped reads (peaks) are called Peak Callers. A detailed description of various ChIP-seq peak callers is presented in (Pepke et al., 2009). Although a number of tools are available, but for this study have used the most efficient and open source tool called Model based Analysis of ChIP-seq (MACS) from (Zhang et al., 2008). Nowadays an upgraded version of MACS called MACS2 is commonly used for this purpose.A MACS2 algorithm does process aligned ChIP-seq bam files both with control and without control samples. Mapped reads are modelled as sequence tags (an integer count of genomic locations mappable under the chosen algorithm). Depending upon the type of protein being ChIPed, different types of peaks are observed when viewing the information in a genome browser, such as Integrated Genome Viewer (IGV). Most of the TFs act as point-source factors and result in narrow peaks, factors such as histone marks generate broader peaks, and proteins such as RNA Pol II can give rise to mixed peaks (both narrow as well as broad). Fig. 3(a) & Fig. 3(b) show the FOXA1 peaks for BRCA1 and ESR1 as viewed in IGV.

**Fig.3:**
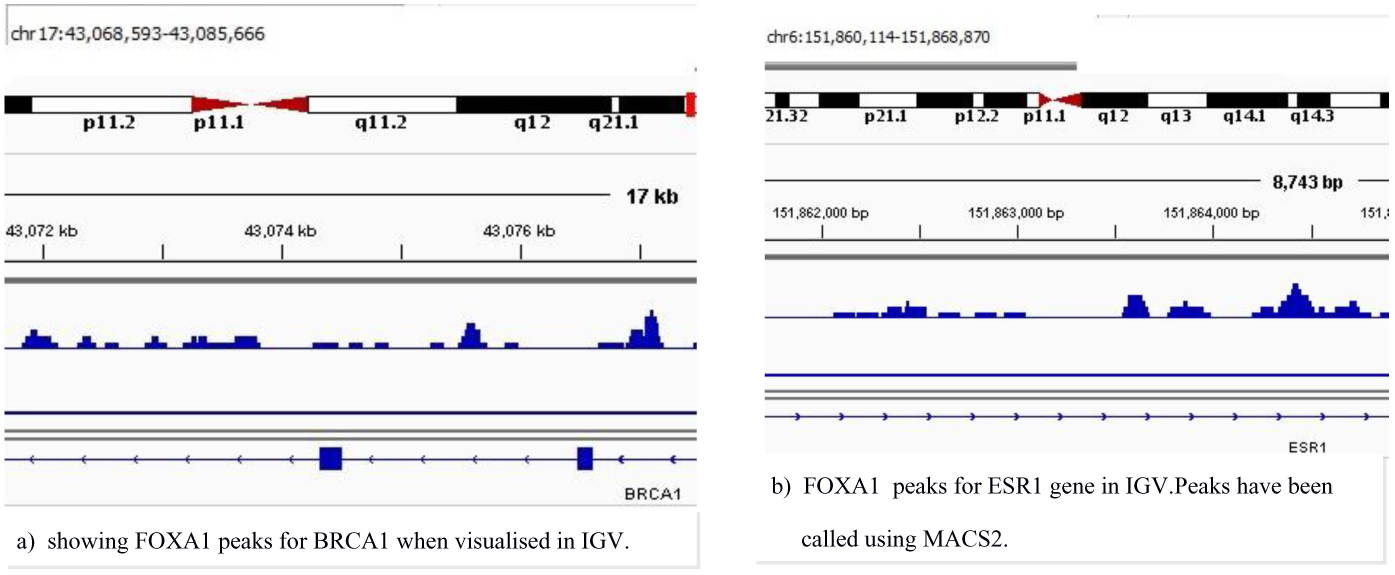
FOXA1 peaks for BRCA1 and ESR1

## 4 Results

The proposed pipeline first generates a list of differentially expressed genes (DEGs) from the normalized expression profiles, thereby identifying the gene activity for both factor-bound and factor-unbound conditions. These DEGs are then integrated with the binding information from stage-III of the pipeline.Both these intermediate data sets are passed as input to stage-IV that employs Binding and expression target analysis (BETA) for prediction process; it calculates binding potential derived from the distance between transcription start site and the TF binding site, thereby modeling the manner in which the expression of genes is being influenced by TF binding sites. Using contributions from the individual TF binding sites, we obtain a cumulative score of overall regulatory potential (probability of a gene being regulated by a factor) of a gene. The percentage of up and down regulated genes is shown in Fig.(4)

**Fig.4:**
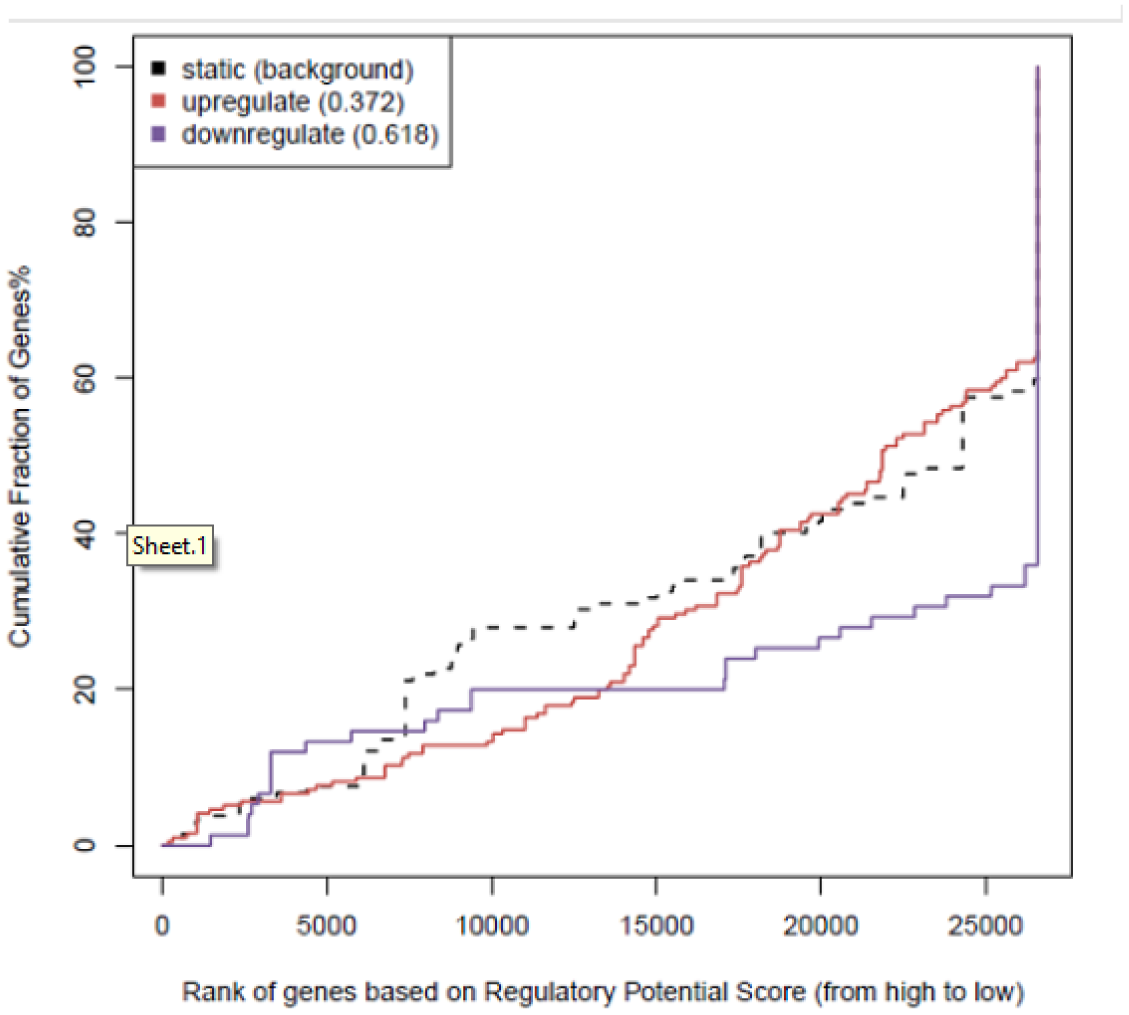
FOXA1 predicted genes activation & repression function

During the process of target prediction each gene receives two ranks, one from the binding potential *R_bp_* and other from differential expression *R_de_*. Both these ranks are multiplied to obtain rank product *R_p_ = R_bp_ × R_de_*. Genes with more regulatory potential and more differential expression are more likely to be as real targets.Table 1 shows a list of up and down regulated genes.

**Table 1:**
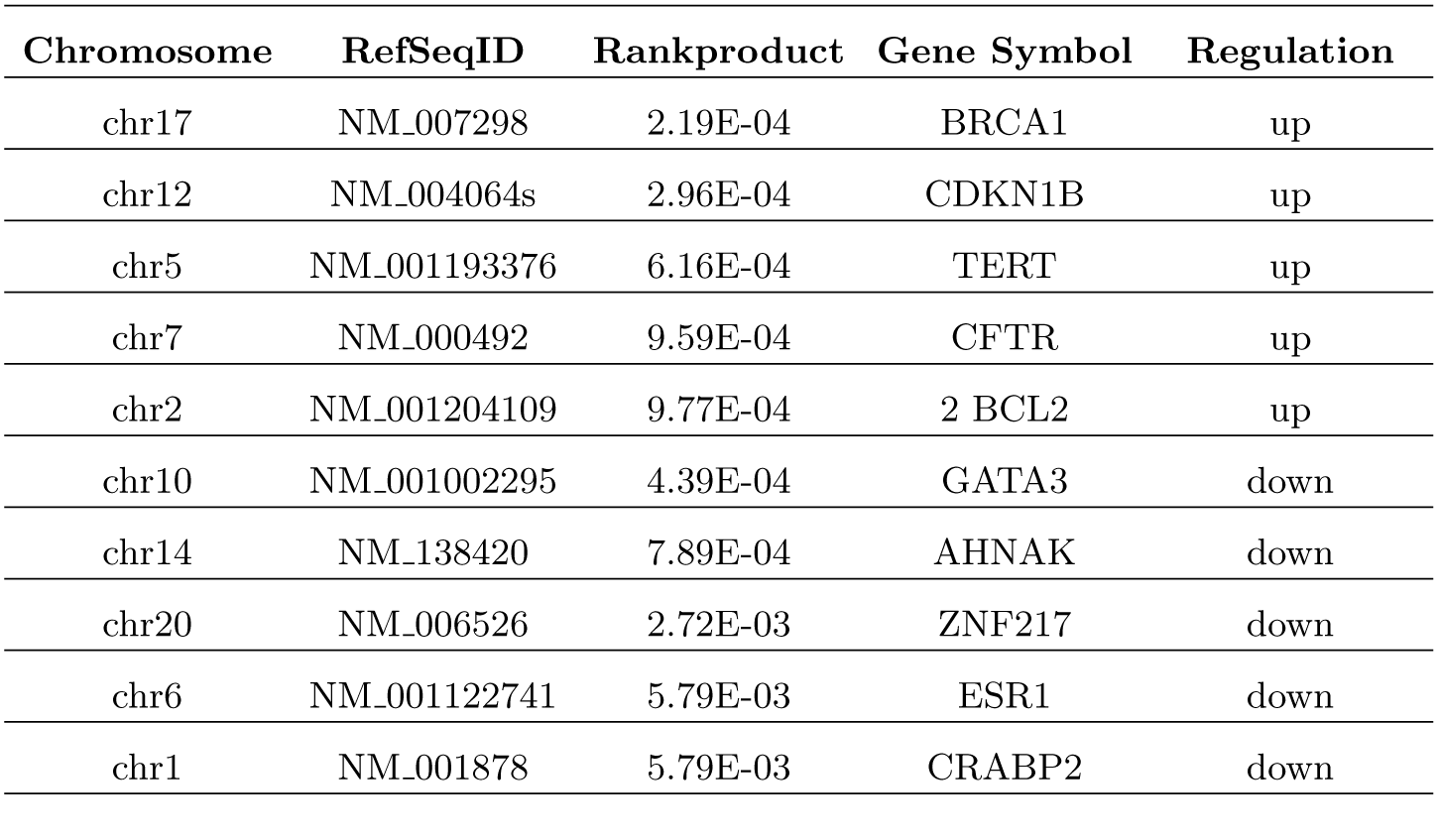
Target genes predicted by inference pipeline

Predicted targets from the inference pipeline results have been widely reported in literature.e.g, FOXA1 up regulated BRCA1 and down regulated ESR1, GATA3 and ZNF217 haven been reported in (Baran-Gale, Purvis, & Sethupathy, 2016).Similarly evidence regarding the role of multiple loci on TERT gene are related to ER(-ve) breast cancer(Bojesen et al., 2013). Many of these predicted targets are well known prognostic biomarkers whose role has been established well in the scientific literature. Once we have a set of target genes, a further downstream analysis of these genes can be done by using gene ontologybased tools such as DAVID to map them with their corresponding biological functions.

## 5 Discussion & Conclusion

The availability and expansion of ChIP-seq and RNA-seq datasets is fuelling an exponential rise in the number of studies being conducted in the area of integrated computational analysis. The motive behind these research endeavors is to address basic questions about how multiple factor binding is related to transcriptional output within in vivo DNA. The proposed inference pipeline is used to decipher the regulatory relationship between TFs that bind to DNA and their corresponding target genes that they influence resulting in their activation/repression.From RNA-seq and ChIP-seq reads, the pipeline generates one file containing differential expression of genes and the other DNA-binding events in the form of peaks. Both these files are integrated in the final stage to yield targets for the TF/TFs whose peaks file was used.

The inference pipeline presented in this paper extracts target genes and hence the regulatory network for a specific TF that has been ChIPred. In case we are required to build a regulatory network for a set of TFs, we need to input new peak files for every new TF in a loop and record the target genes of this TF and its regulatory influence in a separate file. In the current study, we considered only TFs and their influence on gene expression. However, a wider study can include multiple TFs, methylation data, histone marks and polymerase loading to improve the efficiency of the proposed pipeline.

Deciphering the transcriptional regulatory relationships and understanding the elements of regulatory mechanisms that control gene expression is a key research area of regulatory biology. Therefore computational integration of factor binding and other genome-wide data, such as gene expression will be sought after to extract functionally important connections of a working regulatory code.

## Acknowledgment

The author Nisar Wani acknowledges Teacher Fellowship received from University Grants Commission, Ministry of Human Resources Development, Govt. of India vide letter No. F.BNo. 27-(TF-45)/2015 under Faculty Development Programme.

